# Predictive Analysis of Methylation Patterns in Oral Squamous Cell Carcinoma (OSCC) Using Machine Learning

**DOI:** 10.1101/2025.06.19.660641

**Authors:** Debasree Sarkar

## Abstract

**Background:** Oral and oropharyngeal cancers are the most common types of head and neck cancers, with over 90% originating from squamous cells in the mouth and throat. Chronic tobacco and alcohol use, inflammation, viral infections, betel quid chewing, and genetic predisposition are major risk factors for OSCC, which kills over 100,000 patients annually. Epigenetic mechanisms, such as DNA methylation, can silence tumour suppressor genes, contributing to cancer progression and patient outcomes in oral squamous cell carcinoma (OSCC).

**Objective:** This study aimed to predict prominent methylation signatures that can distinguish OSCC from normal cells.

**Methods:** Machine learning algorithms like Support Vector Machine (SVM), Random Forest (RF), and Multilayer Perceptron (MLP) were implemented using R packages and a balanced training dataset consisting of M-values of methylated CpG sites from 46 matched OSCC and normal adjacent tissue samples.

**Results:** MLP model demonstrated the highest accuracy of 92% on the training dataset and 100% on the blind dataset, even with a reduced feature set of just 10 significantly differentially methylated CpG sites.

**Conclusion:** A highly accurate and generalizable machine learning model was developed using the Multi-Layer Perceptron with multiple layers (MLP-ml) algorithm, which achieved an accuracy of 95% on an independent validation dataset of 15 OSCC tumors and 7 non-tumor adjacent tissue samples.

## 1. INTRODUCTION

Oral and oropharyngeal cancers are the two most common types of cancer that develop in the head and neck region, and more than 90% of these cancers originate from the flat, scale-like squamous cells found in the lining of the mouth and throat. After initiation, tumour cells can deeply invade the local structures and lymph nodes of the neck, leading to further distant metastases even into the aero-digestive tract of the patients, all of which increases the chances of potential recurrence of oral cancers [1]. Epigenetic mechanisms that result in dysregulation of gene expression has been found to play a major role in oral squamous cell carcinoma (OSCC) [2], which claims the lives of more than 100,000 patients worldwide every year [3]. Chronic tobacco and alcohol use, which may have a direct impact on epigenetic regulation of gene expression, constitute two major risk factors for OSCC tumorigenesis, along with supplementary factors like chronic inflammation, viral infections (human papillomavirus), betel quid chewing, and genetic predisposition [4,5]. It is, therefore, of critical importance to understand the role of epigenetic alterations, like aberrant DNA methylation, in the initiation and progression of OSCC.

DNA methylation is a key epigenetic modification that can silence tumour suppressor genes, contributing to the development and subsequent progression of different types of cancers [6]. DNA methylation alterations, such as hypermethylation of tumour suppressor genes, are commonly observed in oral squamous cell carcinoma (OSCC), and is known to influence cancer progression and patient outcomes [7]. In addition, increased expression of DNA methyltransferases (DNMTs) is often found in oral cancers, leading to gene inactivation and chromosomal instability [8]. Therefore, targeting DNA methylation through DNMT inhibitors may offer a novel therapeutic strategy against OSCC. In general, it has been observed that global DNA hypomethylation contributes to the process of OSCC tumorigenesis by multiple potential mechanisms including reduction of methylation at DNA repetitive elements leading to chromosomal instability and demethylation of some of the methylation-silenced promoter regions of proto-oncogenes. Furthermore, specific methylation patterns have been associated with tumour differentiation and nodal involvement, leading to differentially methylated regions (DMRs) being recognized as potential biomarkers for early detection and prognosis in oral cancer, with specific genes like DAPK1 and TIMP3 showing significant associations with clinical outcomes [9].

Machine learning algorithms like Support Vector Machine (SVM), Random Forest (RF), and Artificial Neural Network (ANN), might play a crucial role in the identification of prominent methylation signatures that can distinguish OSCC from normal cells by rapid analysis of the high-dimensional datasets typically produced by genome-wide DNA methylation studies. Machine learning (ML) is a branch of artificial intelligence (AI) deeply rooted in applied statistics, building computational models that use inference and pattern recognition instead of explicit sets of rules. ML focuses on developing computer systems that learn from data and progressively improve their predictive performance, and therefore, can be very efficient in detecting patterns embedded in high-dimensional datasets that might not be explicitly defined and discernible by humans. As such machine learning techniques have become fairly popular among biomedical researchers to study methylation patterns associated with various types of cancers [10–12], including OSCC [13–15].

Despite the highest incidence being in Asia, followed by Europe and North America, with a disproportionately high disease burden in Low- and Middle-Income Countries, South America, particularly Brazil, also has high incidence rates of oral and oropharyngeal cancers, which is unfortunately underreported and largely overlooked by the global research community. In this article, machine learning approaches were utilized to predict methylation patterns associated with OSCC in matched OSCC and normal adjacent tissue samples collected from patients at A.C. Camargo Cancer Center, Sao Paulo, Brazil (publicly available dataset GSE234379) [16]. Another publicly available dataset of OSCC tumors and non-tumor adjacent tissues (GSE178216) from patients at Brazilian National Cancer Institute (INCA, Rio de Janeiro, Brazil) was used as an independent validation dataset [17]. Although there are a few studies that have employed machine learning for decoding methylation patterns in diseases like Tuberculosis [18] and Chagas Cardiomyopathy [19], this is the first report of machine learning being used on a cancer methylome dataset from South America.

## 2. MATERIALS AND METHODS

### 2.1. Dataset Description

#### Training Dataset

The dataset GSE234379 was downloaded from the Gene Expression Omnibus (GEO), a public functional genomics data repository available from the National Center for Biotechnology Information, funded by the government of the United States. This dataset consists of DNA methylation data from 46 matched OSCC and normal adjacent tissue samples determined using a genome-wide platform (Illumina Infinium HumanMethylation450 BeadChip).

#### Independent Dataset

The dataset GSE178216, containing genome-wide methylation data from 7 non-tumor adjacent tissues and 15 tumors from OSCC patients using Illumina Infinium HumanMethylation450 BeadChip, was used as the validation dataset.

### 2.2. DNA methylation analysis

DNA methylation analysis was performed using the R Bioconductor package ‘minfi’ (version 1.54.1) [20], wherein Beta values (proportion of methylation at a specific CpG site) and M-values (log-ratio of methylation) for each probe across samples were determined from the raw IDAT files in the dataset, after pre-processing using Noob (normal-exponential out-of-band), a background correction method with dye-bias normalization. Beta values and M-values are two commonly used measures to represent methylation levels, with the caveat that Beta values are more suitable for visualization and clustering, while M-values are better for statistical modeling and differential methylation analysis. This is because M-values have better statistical properties such as more homoscedasticity (homogeneity of variance/variance does not depend on the mean), which also aligns better with assumptions in most machine learning algorithms. Studies have also shown that M-values often lead to better model accuracy and feature selection performance, and machine learning algorithms typically benefit from the unbounded, more Gaussian-like distribution of M-values. In addition, the log-ratio nature of M-values helps highlight subtle but consistent changes, making them more useful for pattern recognition in classification tasks. Hence, a final set of 67 M-values, which were retained after filtering out the rows with missing values or no variation, were chosen as the input dataset for the machine learning algorithms described in the next section.

Packages like ‘limma’ (v. 3.64.1) [21], ‘IlluminaHumanMethylation450kanno.ilmn12.hg19’ (v. 3.21), ‘DMRcate’ (v. 3.4.0) [22], and ‘ChIPseeker’ (v. 1.44.0) [23] were used for Differential Methylation Analysis, annotation and identification of Differentially Methylated Regions (DMRs), and subsequent DMR analysis, comparison, and visualization, respectively.

### 2.3. Machine Learning

The following machine learning algorithms were implemented using the caret (version 6.0-94) package in R [24]: (a) Naïve Bayes (NB), (b) Support Vector Machines with Linear Kernel (SVM-linear), (c) Support Vector Machines with Radial Basis Function Kernel (SVM-radial),(d) Bagged Classification and Regression Trees (treebag), (e) gradient boosting model using decision trees via XGBoost (xgbTree), (f) Random Forest (RF), and (g) Multi-Layer Perceptron, with multiple layers (MLP-ml). Naïve Bayes is a probabilistic classifier based on Bayes’ theorem with the naive assumption that all features are independent and follow a Gaussian distribution. Support Vector Machines (SVMs) are powerful supervised learning algorithms that try to find the optimal hyperplane that best separates data points from different classes by maximizing the margin between them. Bagging stands for Bootstrap Aggregating, and is an ensemble method that creates multiple bootstrap samples (random samples with replacement) from the training dataset. The ‘treebag’ method in caret refers to a bagging ensemble of decision trees, often known as Bagged CART (Classification and Regression Trees). The xgbTree method in the caret package trains a gradient boosting model using decision trees as base learners that is implemented via the XGBoost library, which sequentially builds trees where each new tree attempts to correct errors made by the previous ones. Random Forest is an ensemble of decision trees built using Bagging (Bootstrap Aggregation) and Random Feature Selection, where a large number of trees are built and their predictions are aggregated to produce a more accurate and robust model. A Multi-Layer Perceptron (MLP) is a type of feed-forward artificial neural network and the ‘Multi-Layer Perceptron with Multiple Layers’ method supports multiple hidden layers, using the RSNNS (Stuttgart Neural Network Simulator) backend.

In each of these methods, 80% of the dataset was used as training data for 5-fold cross-validation, while the remaining 20% was used as the blind/test dataset for model evaluation. During 5-fold cross-validation and model evaluation, several threshold-dependent and threshold-independent performance metrics were used [25]. The ‘pROC’ (version 1.18.5) package [26] was used for plotting the Receiver Operating Characteristic (ROC) curves.

## 4. RESULTS

The dataset GSE234379, consisting of genome-wide DNA methylation data from 46 matched oral cavity cancer and normal adjacent tissue samples generated using the Illumina Infinium HumanMethylation450 BeadChip (450k), was downloaded from the Gene Expression Omnibus (GEO) database, and analyzed using various R packages. Finally, a carefully filtered set of 67 M-values (representing 67 CpG sites) was used as input to train machine learning models using NB, SVM-linear, SVM-radial, treebag, xgbTree, RF, and MLP-ml algorithms, as described in the Materials and Methods section. As shown in **Table 1A & B**, MLP-ml model produced the best accuracy score of 92% on the training set, and 100% on the blind dataset.

To further analyze the MLP-ml model, the number of features (CpG sites) used for prediction were sequentially reduced to determine the least number of features sufficient to develop a minimalistic model. As shown in **Table 2**, the best performing minimalistic model used only the top 10 features to give an accuracy score of 100%. **Table S1** lists the details of these top 10 CpG sites including the summary test statistic for the DMR(from limma), mean difference in M-values across the DMR, p-value for the DMR before any correction for multiple testing, and FDR-adjusted p-value using the Benjamini-Hochberg method, along with the genomic co-ordinates and overlapping genes (if any). Interestingly, the first two CpG sites in the list correspond to the genes CCDC17 and SELI/SELENOI, which have already been implicated in various cancers [27,28], including squamous cell carcinoma (SCC) [29].

**Table 1A:** Summary of performance metrics for different machine learning methods on the training dataset.

|  | logLoss | AUC | F1 | Sensitivity | Specificity | Precision | Recall | Accuracy |
| --- | --- | --- | --- | --- | --- | --- | --- | --- |
| NB | 2.42 | 0.89 | 0.85 | 0.86 | 0.84 | 0.84 | 0.86 | 0.85 |
| svmL | 0.40 | 0.89 | 0.84 | 0.84 | 0.84 | 0.84 | 0.84 | 0.84 |
| svmR | 0.41 | 0.90 | 0.88 | 0.89 | 0.86 | 0.87 | 0.89 | 0.88 |
| treebag | 0.28 | 0.95 | 0.90 | 0.95 | 0.84 | 0.85 | 0.95 | 0.89 |
| xgbTree | 0.30 | 0.97 | 0.91 | 0.95 | 0.86 | 0.88 | 0.95 | 0.91 |
| RF | 0.29 | 0.97 | 0.91 | 0.95 | 0.86 | 0.88 | 0.95 | 0.91 |
| mlpML | 0.30 | 0.94 | 0.92 | 0.95 | 0.89 | 0.90 | 0.95 | 0.92 |

**Table 1B:**
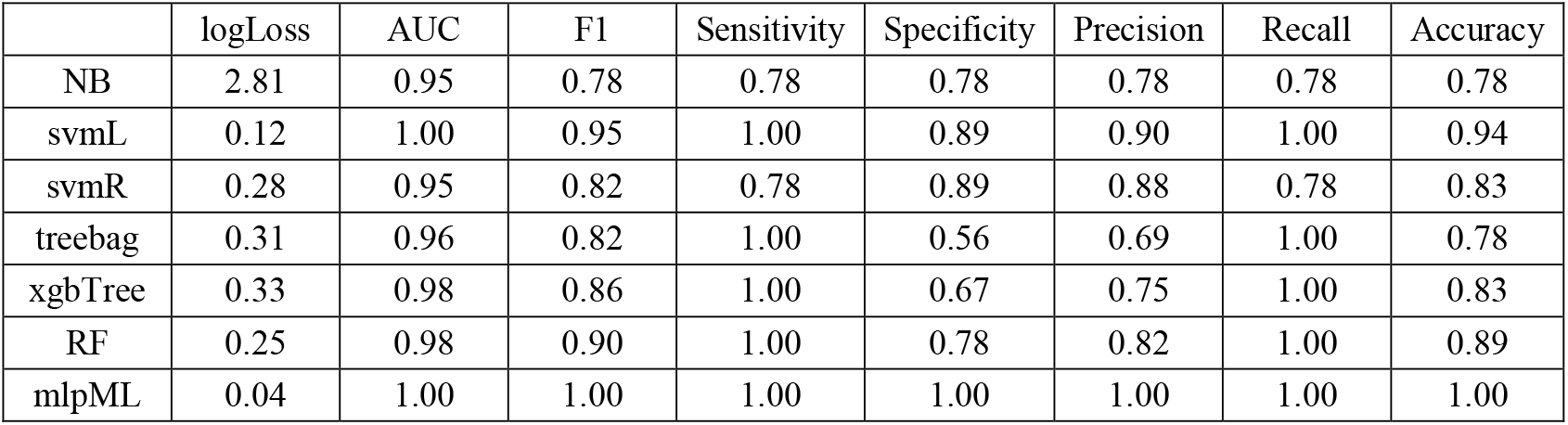
Summary of performance metrics for different machine learning methods on the blind dataset.

**Table 2:**
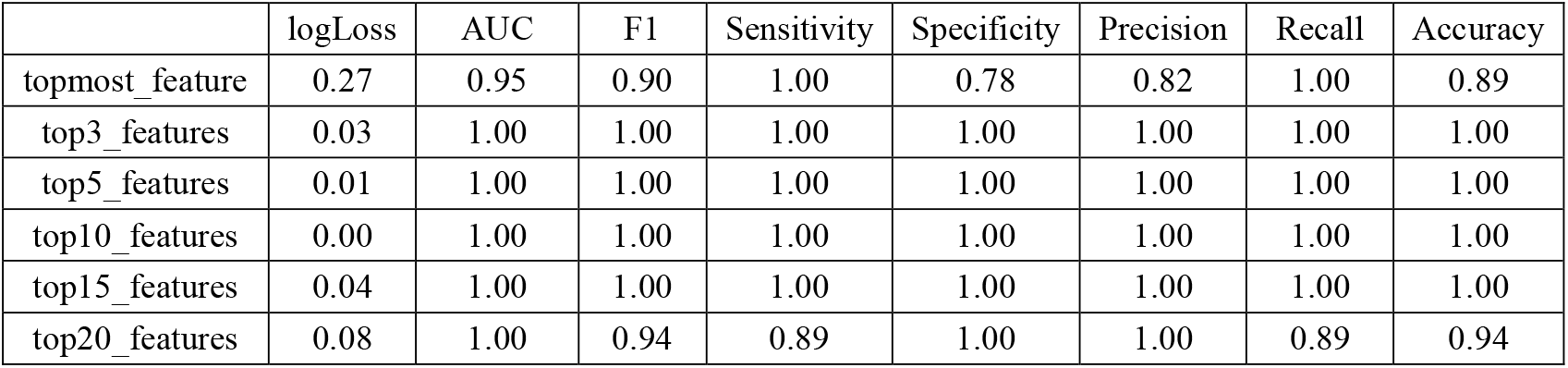
Summary of performance metrics for mlpML using different number of features.

The minimalistic MLP-ml model using only the top 10 CpG sites as features was also used to predict OSCC samples in an independent validation dataset GSE178216 consisting of 15 OSCC and 7 normal adjacent tissue samples, achieving Accuracy of 0.9545 [95% CI : (0.7716, 0.9988), P-Value 0.002469] and AUC of 100%, as shown in Table 3.

**Table 3:** Summary of performance metrics for the minimalistic MLP-ml model using only the top 10 CpG sites as features on the independent validation dataset.

|  | logLoss | AUC | F1 | Sensitivity | Specificity | Precision | Recall | Accuracy |
| --- | --- | --- | --- | --- | --- | --- | --- | --- |
| GSE178216 | 0.11 | 1.00 | 0.93 | 1.00 | 0.93 | 0.88 | 1.00 | 0.95 |

## 5. DISCUSSION

The incidence of oral cancer and corresponding mortality rates in South America is notably high [30], with Brazil reporting the highest rates among males [31]. The rising incidence rates, driven by factors such as tobacco and alcohol consumption, poses significant challenges for public health policy, necessitating targeted interventions to address the underlying risk factors and improve healthcare access. In the current study, 2 whole-genome methylation datasets originating from Brazilian hospitals were analyzed using machine learning algorithms to decipher specific methylation patterns associated with OSCC. The M-value (log-ratio of methylation for each probe) matrix of 485512 probes across 92 samples (46 OSCC tumour & 46 matched normal) were screened to remove rows with missing values or no variation, yielding a curated set of 67 CpG sites. This is a crucial pre-processing step because rows with NAs (missing values) can corrupt statistical integrity or break functions, while imputation of methylation values is tricky and can introduce bias, especially for high-dimensional, sparse data like methylation arrays. Removal of rows with zero variance, on the other hand, is intuitive because methylation sites with the same value in all samples cannot distinguish between the sample groups, but add noise or redundancy, increasing model complexity without any benefit.No-variation sites are biologically uninformative and computationally inefficient, they inflate dimensionality without contributing to variance, possibly distorting results, and hence should be excluded.

Machine learning models from several methods including NB, SVM-linear, SVM-radial, treebag, xgbTree, RF, and MLP-ml algorithms were trained using the larger dataset (GSE234379), which is a balanced dataset of 46 OSCC and 46 normal samples. Use of a balanced training set in machine learning is crucial to prevent the model from overfitting, and allow better generalization, more reliable performance metrics, and more informative feature importance rankings. The generalization ability of the different machine learning models was then checked using the second Brazilian dataset (GSE178216), which was generated using the same Illumina Infinium HumanMethylation450 BeadChip, as an independent dataset. The MLP-ml model achieved the best prediction accuracy on the training examples, as well as on the independent dataset, even with a reduced feature set of only top 10 CpG sites. MLP models have been previously used to predict driver genes from multi-omics pan-cancer data, which included DNA methylation profiles [32], as well as risk of diabetes and cancer from DNA methylation arrays [33].

## CONCLUSION AND FUTURE DIRECTIONS

Overall, this study provided clues into salient methylation signatures unique to OSCC in South American, or more specifically Brazilian patients, from two publicly available whole genome methylation datasets using machine learning prediction models. The best performing minimalistic MLP model used only the top 10 CpG sites to give an accuracy score of 100% on both the blind testing set as well as the second independent validation dataset. Similar studies are needed on methylation datasets from other South American countries, for further validation of our model and the methylation pattern associated with OSCC in our study. In addition, this study focuses only on DNA methylation, but a more comprehensive model should also incorporate other epigenetic signals, as well as, correlate them with gene expression datasets, for a holistic overview of the underlying mechanistic aspects of OSCC tumorigenesis and disease progression.

## SUPPLEMENTARY INFORMATION

**TABLE S1:**

| DMR | Statistical parameters |  |  |  | Genomic Co-ordinates and Overlapping Gene Information |  |  |  |  |  |
| --- | --- | --- | --- | --- | --- | --- | --- | --- | --- | --- |
|  | stat | diff | rawpval | fdr | chr | pos | strand | UCSC_RefGene_Name | UCSC_RefGene_Accession | UCSC_RefGene_Group |
| cg00702837 | -7.8553132 | -0.2392556 | 4.41E-12 | 1.48E-10 | chr1 | 46090494 | - | CCDC17 | NM_001114938 | TSS1500 |
| cg19045284 | -5.3743876 | -0.2374391 | 4.95E-07 | 2.21E-06 | chr2 | 26572101 | + | SELI | NM_033505 | Body |
| cg09991426 | -7.603319 | -0.2511719 | 1.53E-11 | 3.41E-10 | chr2 | 83316213 | + | — | — | — |
| cg06672958 | -10.120889 | -0.2268153 | 4.85E-17 | 3.25E-15 | chr5 | 106830972 | + | EFNA5 | NM_001962 | Body |
| cg10093438 | -7.0258164 | -0.2315154 | 2.54E-10 | 3.40E-09 | chr7 | 138089004 | - | — | — | — |
| cg11523381 | -6.3814464 | -0.1777731 | 5.37E-09 | 4.50E-08 | chr11 | 63751153 | - | — | — | — |
| cg18275339 | -6.1698024 | -0.2855646 | 1.43E-08 | 1.06E-07 | chr11 | 91760338 | + | — | — | — |
| cg26199225 | -7.0466065 | -0.1897635 | 2.30E-10 | 3.40E-09 | chr14 | 100673716 | - | — | — | — |
| cg21387117 | -6.8195749 | -0.1653235 | 6.82E-10 | 6.52E-09 | chr17 | 3288348 | - | — | — | — |
| cg09614762 | -6.8631874 | -0.176285 | 5.54E-10 | 6.18E-09 | chrX | 65038012 | - | — | — | — |

stat represents the mean or combined test statistic for all CpGs in the region, often derived from moderated t-statistics from the underlying linear modelling;
diff represents the average methylation difference between the two groups, where Positive values → Hypermethylation & Negative values → Hypomethylation;
rawpval represents the original/uncorrected p-value for the DMR before any correction for multiple testing, where lower values indicate stronger evidence of differential methylation;
fdr represents the False Discovery Rate (FDR)-adjusted p-value using the Benjamini-Hochberg method, used to assess statistical significance of DMRs while controlling for multiple comparisons;
chr is the chromosome number; pos is the genomic location.

**Figure S1:**
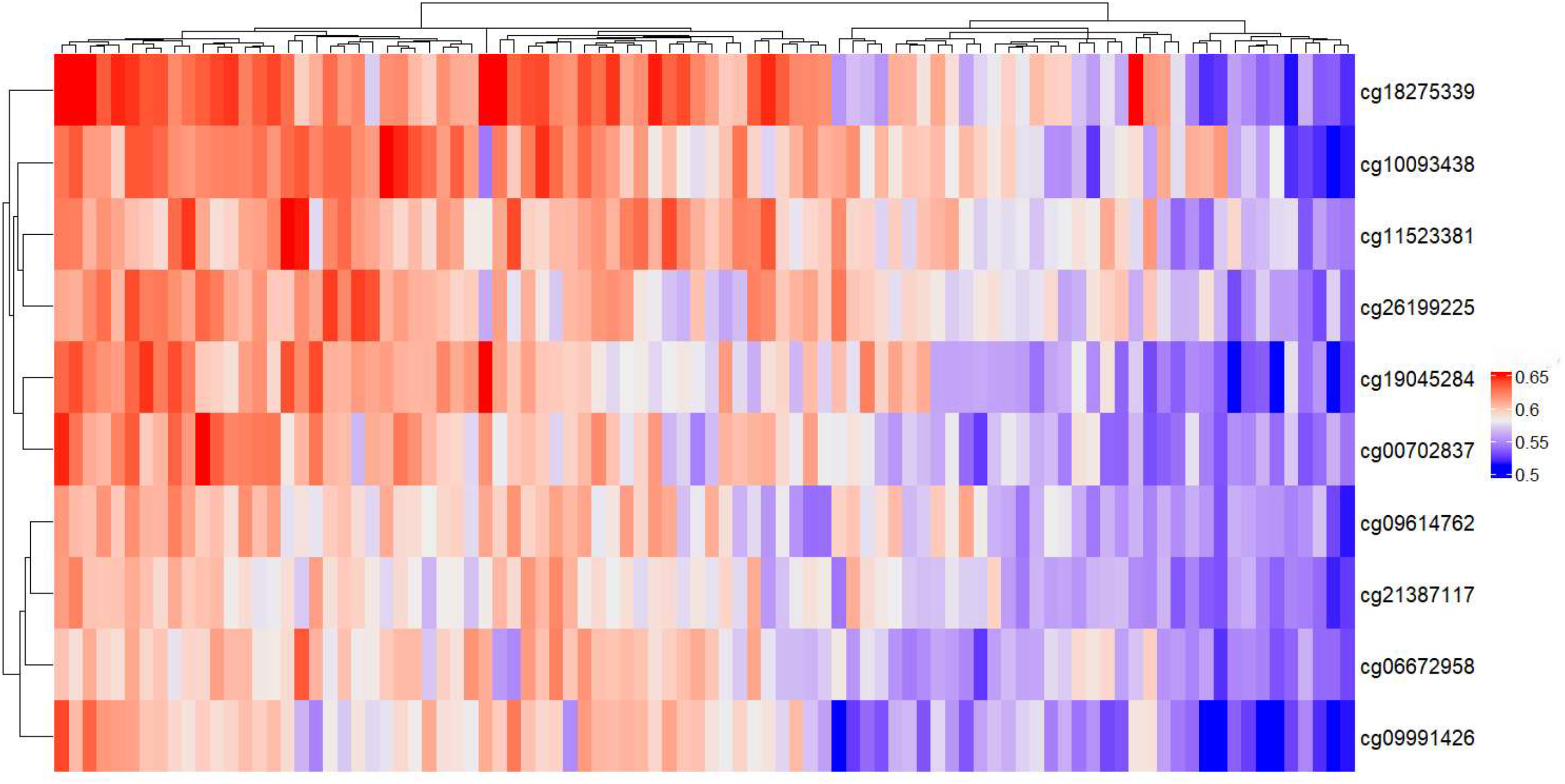
Heatmap of Beta values of the top 10 CpG sites detected by machine learning in GSE234379 samples (N=92: 46 OSCC & 46 matched adjacent normal tissue samples).

**Figure S2:**
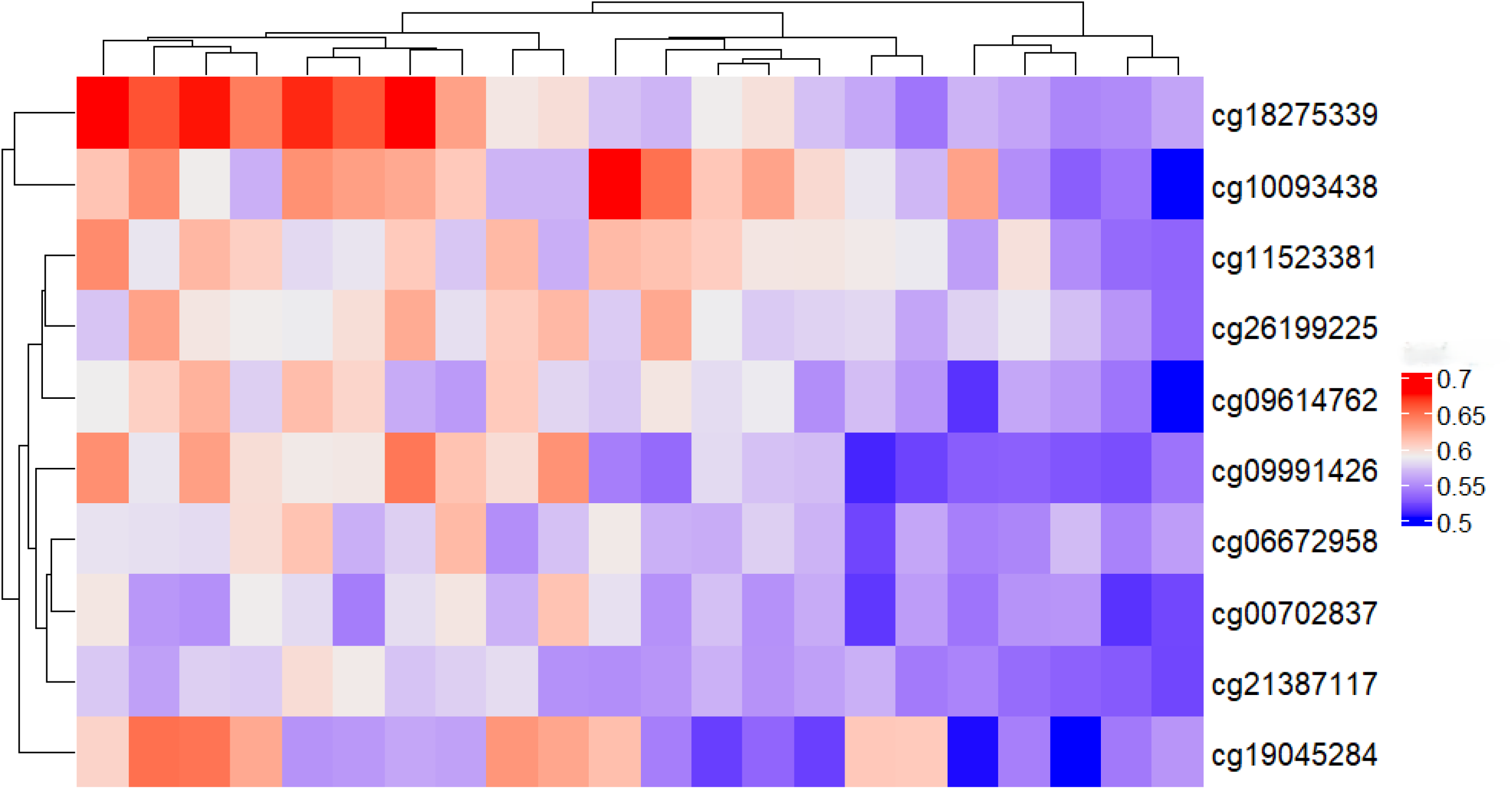
Heatmap of Beta values of the top 10 CpG sites detected by machine learning in GSE178216 samples (N=22: 15 OSCC and 7 normal adjacent tissue samples).

**Figure S3:**
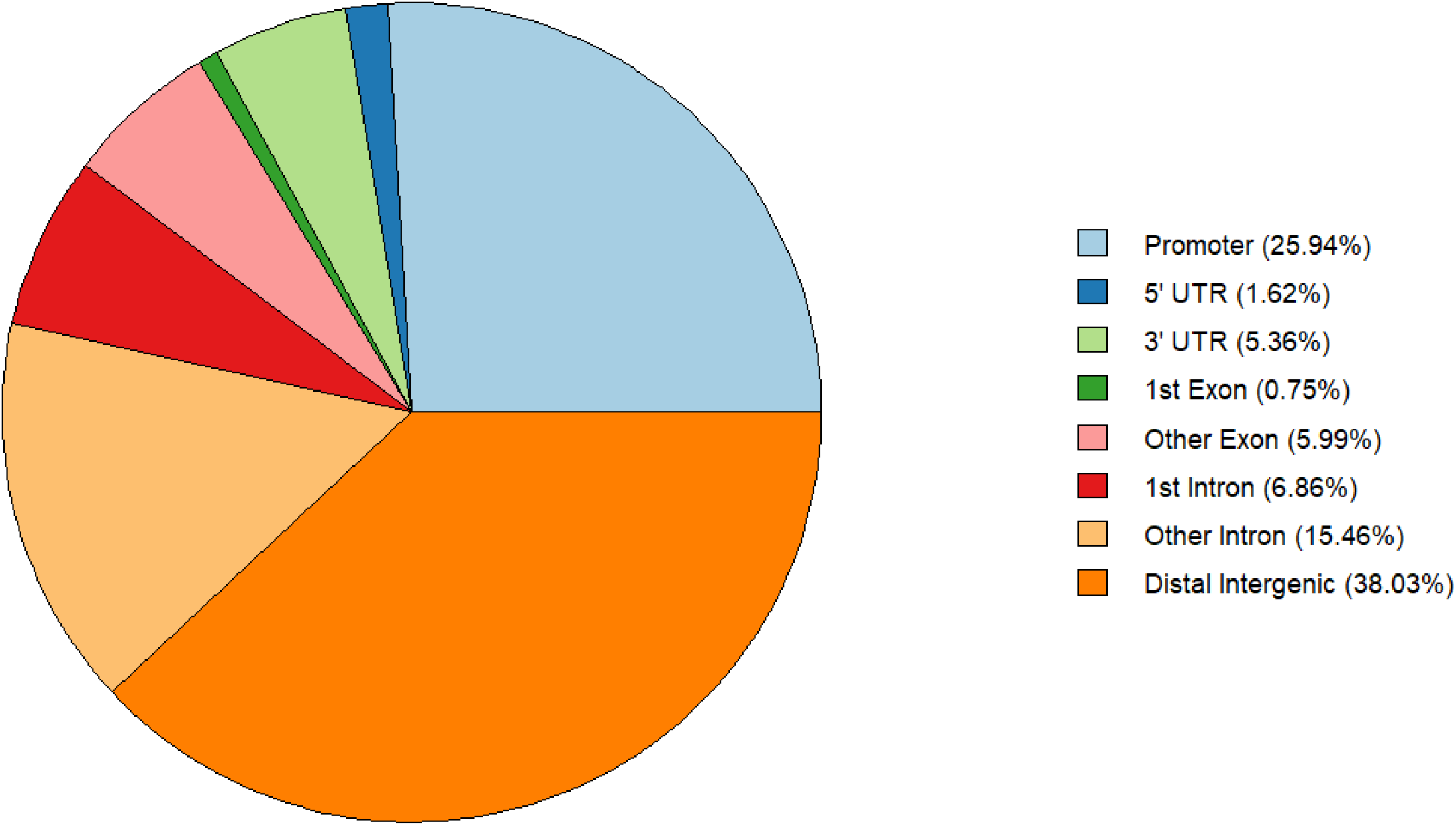
Pie chart of all annotated differentially methylated regions in GSE234379 obtained using ‘limma’, ‘DMRcate’, and ‘ChIPseeker’ packages.

